# Identification of the Molecular Basis of Anti-fibrotic Effects of Soluble Guanylate Cyclase Activator Using the Human Lung Fibroblast Phosphoproteome

**DOI:** 10.1101/2020.06.12.148908

**Authors:** Sunhwa Kim, Ashmita Saigal, Weilong Zhao, Peyvand Amini, Alex M. Tamburino, Subharekha Raghavan, Maarten Hoek

## Abstract

Idiopathic pulmonary fibrosis (IPF) is an irreversible and progressive fibrotic lung disease. Advanced IPF patients often demonstrate pulmonary hypertension, which severely impairs patients’ quality of life. The critical physiological roles of soluble guanylate cyclase (sGC)-cyclic guanosine monophosphate (cGMP) pathway have been well characterized in vasodilation and the corresponding therapies and pathway agonists have shown clinical benefits in treating hypertension. In recent years, many preclinical studies have demonstrated anti-fibrotic efficacy of sGC-cGMP activation in various experimental fibrosis models but the molecular basis of the efficacy in these models are not well understood. Also, sGC pathway agonism has demonstrated encouraging clinical benefits in advanced IPF patients (NCT00517933). Here, we have revealed the novel phosphorylation events downstream of sGC activation in human lung fibroblasts using phosphoproteomics. sGCact A, a potent and selective sGC activator, significantly attenuated more than 2,000 phosphorylation sites. About 20% of phosphorylation events, attenuated by transforming growth factor β (TGFβ), a master regulator of fibrosis, were further dysregulated in the sGCact A co-treated lung fibroblasts. The overall magnitude and diversity of the sGCact A phosphoproteome was extensive. Further investigation would be required to understand how these newly identified changes facilitate human pulmonary fibrosis.

## Introduction

Idiopathic pulmonary fibrosis (IPF) is an irreversible fibrotic lung disease with unknown etiology [1–3]. Although two approved medications, pirfenidone and nintedanib, are able to slow down lung function decline in IPF patients, many other chronic pathologic conditions such as dyspnea and pulmonary hypertension (PH) and overall disease progression, as measured by progression free survival and established lung fibrosis are not well managed [4]. The median survival rate of IPF patients is 3 to 5 years from diagnosis and particularly, IPF-PH patients show a 3-fold increase in mortality as compared to IPF populations without PH [5, 6].

For more than a century, nitrates have been used to treat PH as a vasodilator. However, exogenous NO donors turned out to be limited by increased oxidative stress and tolerance. Thus, current treatment strategies focus on inhibiting the degradation of cGMP by targeting phosphodiesterases (PDE), mainly PDE type 5 (PDE5) and the generation of cGMP by increasing the enzymatic activity of sGC, mainly by sGC activators (sGCact) and stimulators (sGCstim), which agonize the activity of oxidized and reduced forms of sGC enzyme, respectively.

Preclinical benefits of sGC-cGMP pathway activation have been explored in various disease model settings with or without vascular disorders. Accumulated evidences have highlighted novel benefits of the sGC pathway activation in fibrosis and/or inflammation. Both sGCact and sGCstim significantly ameliorated diet- and/or injury-induced steatosis, inflammation, and/or fibrosis in experimental fibrosis models in the liver [7–9], skin [10], and/or kidney [11]. For example, an sGCact, BAY60-2270, significantly slowed down fibrosis progression in a CCl4-induced liver fibrosis model [7]. Both the level of hydroxyproline, a major component of the collagens (COL) and the areas of fibrosis in liver were significantly lowered. BAY41-2272, an sGCstim significantly delayed bleomycin-induced skin fibrosis by decreasing skin thickness and lowering hydroxyproline levels [10]. Another sGCstim, praliciguat/ IW-1973 also significantly reduced high fat diet-induced steatosis, inflammation and fibrosis [8]. These preclinical findings in various organ systems with different insults of injuries strongly suggest a general anti-fibrotic effect of sGC agonism. Indeed, sildenafil, a PDE5 inhibitor, improved diffusing capacity for carbon monoxide test, oxygenation and St. George’s Respiratory Questionnaire total score in IPF patients, although no statistically significant benefit was measured in the 6-minute walk distance test [12].

Despite the broad understanding of the molecular basis of the sGC pathway activation in vasodilation, primarily mediated through protein kinase G (PKG) and the activation of its down-stream signaling cascades [13–15], the mechanism of how sGC pathway activation can deliver anti-fibrotic effects is not well defined. Small-scale western blot analyses have identified TGFβ-activated extracellular signal-regulated kinase (ERK) as a target of sGC activator BAY41-2272 or cGMP analogue 8-Bromo-cGMP in primary human and rodent dermal fibroblasts [16].

In the present study, we applied phosphoproteomic technology to unbiasedly obtain broad understanding of the overall sGC activation-induced phosphorylation events in human lung fibroblasts. The singleton effect of a sGC activator and its effect on TGFβ signaling are evaluated and discussed.

## Materials and Methods

### Cell culture and stimulation

HFL1 (human lung fibroblasts) was purchased from ATCC (CCL-153) and grown up to 10 passages in complete culture media of F12K media (ATCC, 30-2004) containing 10% v/v fetal bovine serum (ATCC, 30-2020). Recombinant human TGFβ was purchased from R&D (240-B-010). TGFβ was incubated with any cells for 30 minutes to 48 hours.

### Compound preparation

sGCact A (US Patent Number 8455638 B2) and sGC stimulator (US Patent Number 9365574 B2) were synthesized at Merck & Co., Inc., Kenilworth, NJ, USA. TGFβR inhibitor (SB-525334) was purchased from Sigma Aldrich. All compounds were dissolved in dimethyl sulfoxide (DMSO) to stock concentrations of 10 or 1 mM. The final concentration of DMSO in all experiments did not exceed 0.1% v/v.

### Phosphoproteome sample preparation, liquid chromatography (LC), mass spectrometry (MS)

HFL1 cells were used for the experiments. Prior to the experiments, HFL1 cells were cultured in complete culture media for 24 hours and then serum starved for overnight. Post 30 minutes with sGCact A or SB-525334 (10 uM or 1 uM of final concentration, respectively), HFL1 cells were stimulated with recombinant TGFβ at 200 ng/ml (final concentration) for 30 minutes. The cells were quickly washed with ice cold phosphate-buffered saline (Thermo Fisher Scientific, 10010023). The LC-MS analyses, data processing and analyses were conducted at Evotec with their standard procedures [17–20].

### Ingenuity pathway analyses (IPA)

The enriched phosphopeptides, selected by p value and/or fold change (FC) as indicated were uploaded into the IPA software (Qiagen). The Core Analyses function included in the software was used to interpret the data for top canonical pathways.

### Compound profiling in BioMAP^®^ diversity plus and fibrosis systems

The study was conducted at DiscoverX, a part of Eurofins with their standard procedures. Briefly, 15 human primary cell-based systems were pre-treated with sGC stimulator or activator for 30 minutes prior to the incubation with dynamic stimuli for additional 24-48 hours. These 15 systems designed to model different aspects of the human body in an in vitro format. Tissue fibrosis biology is modeled in a pulmonary (SAEMyoF system) and a renal (REMyoF) inflammation environment, as well as in a simple lung fibroblasts (MyoF). Vascular biology is modeled in both a Th1 (3C system) and a Th2 (4H system) inflammation environment, as well as in a Th1 inflammatory state specific to arterial smooth muscle cells (CASM3C system). Additional systems recapitulate aspects of the systemic immune response including monocyte-driven Th1 inflammation (LPS system) or T cell stimulation (SAg system), chronic Th1 inflammation driven by macrophage activation (*/*Mphg system) and the T cell-dependent activation of B cells that occurs in germinal centers (BT system). The BE3C system (Th1) and the BF4T system (Th2) represent airway inflammation of the lung, while the MyoF system models myofibroblast-lung tissue remodeling. Skin biology is addressed in the KF3CT system modeling Th1 cutaneous inflammation and the HDF3CGF system modeling wound healing.

### Homogeneous time resolved fluorescence (HTRF) analyses

The treated HFL lung fibroblasts were lysed for analysis of SMAD3 pSer423/425 (cat#63ADK025PEH, cat#64ND3PEH; CisBio) and VASP pSer239 (cat#63ADK065PEG, cat#63ADK067PEG; CisBio) and the phosphorylation signals were normalized to the total level of SMAD3 or VASP, respectively. The assays were performed as per manufacturer’s instruction and the signal was captured using an Envision plate reader.

### RNA isolation and quantitative real-time PCR (qPCR) analysis

RNA was isolated with the RNeasy mini kit (Qiagen) according to the manufacturer’s instructions. Synthesis of complementary DNA, real-time PCR followed the manufacturer’s instructions. All gene expression results are expressed as fold change relative to the housekeeping genes α-actin, glyceraldehyde 3-phosphate dehydrogenase (gapdh) and/or hypoxanthine-guanine phosphoribosyltransferase (hgprt). TaqMan probes were purchased from Life Technologies.

### Statistical analyses

Differences of each treatments were tested for their statistical significance by Mann– Whitney U non-parametric test unless otherwise indicated. p values equal or less than 0.05 were considered significant.

## Results

### sGC Activator Significantly Attenuated Inflammation and Fibrosis

To broadly investigate the biological effects of sGC pathway activation, two selective and potent agonists of the sGC enzyme, sGC activator (sGCact A: US patent No. 8455638 B2) and sGC stimulator (US patent No. 9365574 B2), were profiled in primary human cells systems that recapitulate key aspects of various human diseases and pathologic conditions by co-culturing distinct population of cells with dynamic repertoire of stimulation (BioMAP^®^ diversity plus and fibrosis panels).

sGCact A significantly attenuated several pathologic aspects in the pulmonary fibrosis system, which recapitulates pathologic features of chronic inflammation and fibrosis by co-culturing human small airway epithelial cells and lung fibroblasts along with the stimulation of both TGFβ and TNFα (tumor necrosis factor alpha). sGCact A significantly and dose-dependently inhibited α-smooth muscle actin (SMA), COL III and interferon-inducible T cell alpha chemoattractant (ITAC), measuring myofibroblast activation, fibrosis related matrix activities and inflammation-related activities, respectively (Fig 1). In contrast to the pulmonary fibrosis system, the effect of sGCact A in the renal fibrosis system, which contained renal proximal tubule epithelial cells and fibroblasts, only showed decreased matrix metalloproteinase 9 expression (Suppl Fig 1A). In the BioMAP^®^ diversity panel, sGCact A significantly decreased soluble (s)TNFα and vascular cell adhesion protein (VCAM)-1, measuring inflammation-related activities and plasminogen activator inhibitor (PAI)-1 and tissue inhibitor of metalloproteinases (TIMP)2, measuring tissue remodeling activities in the systems of chronic inflammation and/or autoimmunity conditions (Suppl Fig 1B). The sGC stimulator also significantly decreased the expression of COL III, αSMA, monocyte chemoattractant protein (MCP)1, and VCAM-1 in renal or pulmonary fibrosis or chronic inflammation conditions (Suppl Fig 1C). Since the overall effect of sGCact A was more active in the pulmonary fibrosis system as compared to the effect of sGC stimulator, we further continued our studies with sGCact A to investigate its effects in lung fibroblasts.

**Fig 1.**
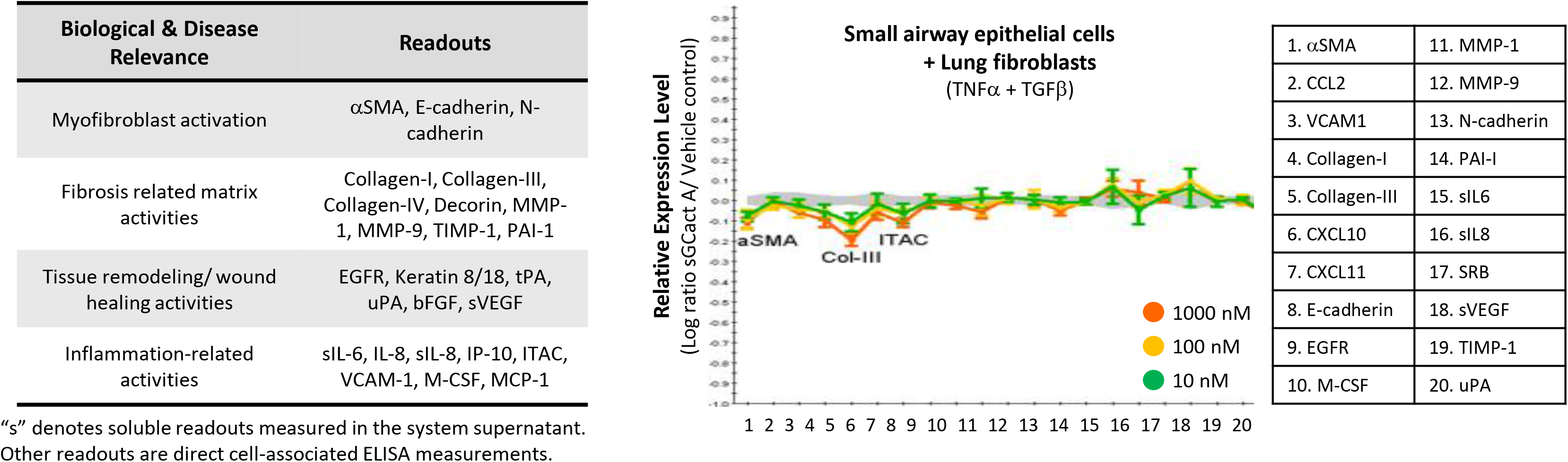
sGC activation-mediated inhibition of fibrosis in human cells. Profiling of sGCact A in BioMAP^®^ Fibrosis panel. X-axes list the quantitative protein marker readouts. Y-axes show log-transformed ratios of readouts. The grey region around the Y-axis represents the 95% significance envelope generated from historical vehicle controls. Biomarker activities are anointed when 2 or more consecutive concentrations change in the same direction relative to vehicle controls, are outside of the significance envelope, and have at least one concentration with an effect size > 20%.

### sGC Activator-induced Changes in Cellular Phosphoproteomics

sGC agonism decreased ERK phosphorylation in TGFβ-treated dermal fibroblasts. To broaden our understanding of sGC agonism-induced molecular changes, we measured phosphopeptides in the human lung fibroblasts, pretreated with sGCact A and then stimulated with TGFβ. We also pretreated cells with SB-525334, a small molecule compound which abrogates both canonical and non-canonical pathways of TGFβ and inhibits TGFβ-induced fibrosis. The changes from sGCact A and SB-525334 were compared to elucidate any similarities.

The treated HFL1 cells were processed and analyzed by liquid chromatography-mass spectrometry (LC-MS) and MaxQuant program to identify sGCact A-induced changes in phosphopeptides (Fig 2A). The activities of sGCact A, TGFβ and SB-525334 in lung fibroblasts were assessed prior to LC-MS. TGFβ-induced phosphorylation at Ser423/425 of Smad3 (pSmad3), a primary transcription factor in the canonical TGFβ pathway [21, 22], was measured by the HTRF assay (Suppl Fig 2A). SB-525334 decreased TGFβ-induced pSmad to the basal level (p<0.01 as compared to the TGFβ-treated sample). No change was measured in the sGCact-treated cells. In a separate experiment, we confirmed little to no inhibitory effect of sGCact A on TGFβ-induced pSmad2-Ser^465/467^ and pSmad3-Ser^423/425^ (Suppl Fig 2B). The sGCact A-treatment mildly increased pVASP-Ser^157^ (p<0.01 as compared to the vehicle-treated sample), whose phosphorylation was markedly elevated by riociguat, an approved sGC stimulator, in human platelets [23] (Suppl Fig 2C). The elevation of pVASP-Ser^157^ upon treatment with sGC stimulator was equivalent to the sGCact A effect (data not shown). There was little to no change in VASP phosphorylation events upon TGFβ stimulation with or without SB-525334 (Suppl Fig 2C).

**Fig 2.**
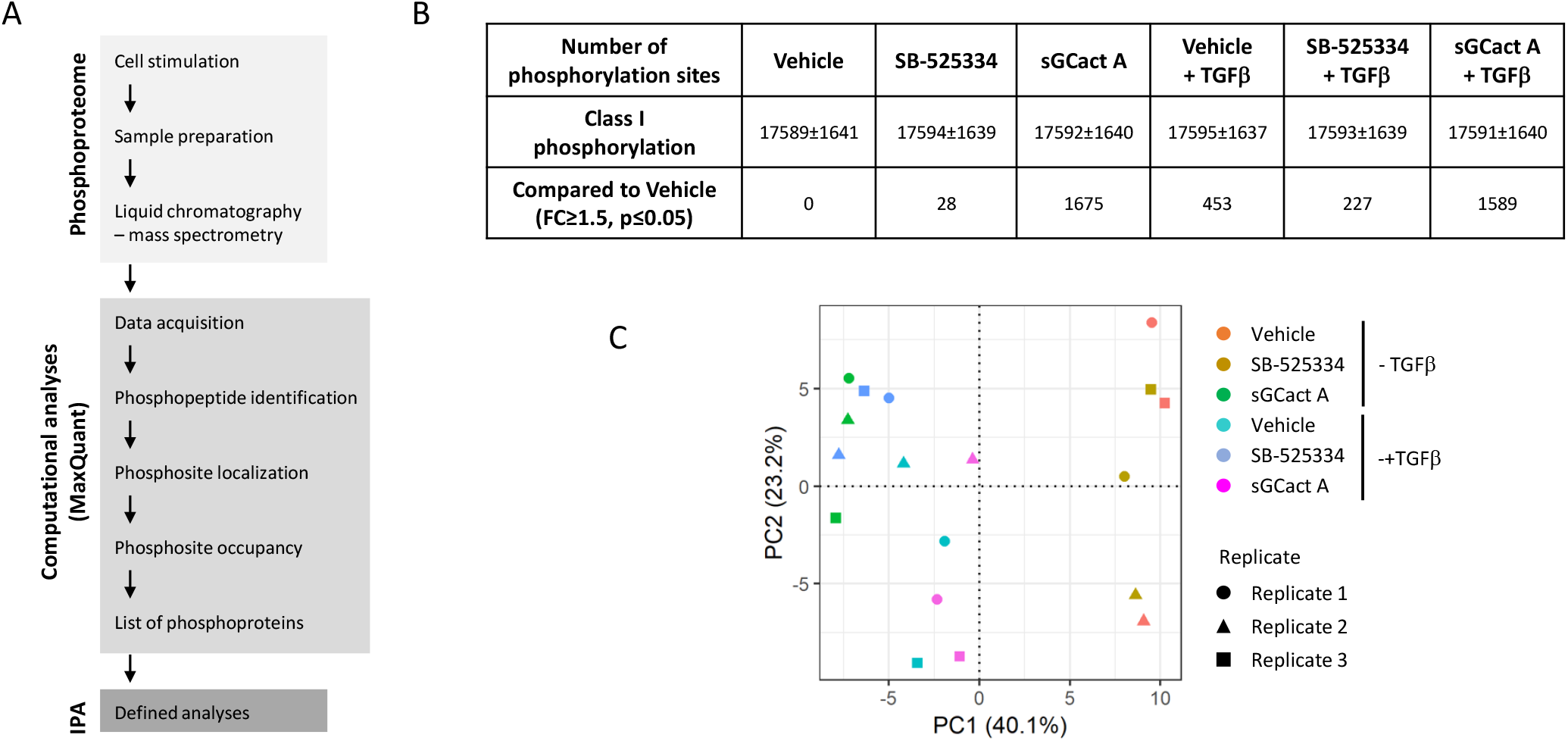
Identification of phosphopeptides, regulated by the sGC activator (sGCact A) in human lung fibroblasts. (A) Workflow for phosphoproteome study. Three independent samples were prepared and processed separately. Post the phosphopeptide identification (FDR at 1%), the data were combined and analyzed at IPA. (B) Quantified class I phosphorylation sites across the replicates and the conditions. More than 19,000 class I phosphorylation sites were identified. (C) PCA of the acquired data virtual pool normalized intensities. To eliminate the batch effects, the data were normalized to a virtual pool as described at Methods.

All treatments quantified similar number of total phosphopeptides with a mean of approximately 17,600 (Fig 2B). A total of 23,885 unique phosphosites were measured from the combined datasets (Suppl Excel 1). Each treatment significantly attenuated multiple phosphopeptides as compared to those in the vehicle-treated samples (fold change (FC)≥1.5, p≤0.05; Fig 2B). Almost a 7-fold increase in phosphopeptides were identified from the sGCact A-treated cells (alone or with TGFβ) compared to the cells treated with SB-525334 (alone or with TGFβ) (1632±61 vs 236±213) (Fig 2B). PCA separated and identified two clusters, either treated with sGCact A or TGFβ or none (Fig 2C).

### Dynamic Changes of TGFβ-induced Phosphorylation Events by SB-525334

TGFβ stimulation induced dynamic changes in phosphorylation events as shown in the heat map signature (Fig 3A). Overall, TGFβ treatment attenuated a total of 453 phosphopeptide, with 215 up-regulated and 238 down-regulated as compared to vehicle-treated fibroblasts (FC≥1.5, p≤0.05, Fig 3B and Suppl Excel 2). As expected, SB-525334 antagonized many TGFβ-induced phosphorylation changes (Figs 3A and 3B). Of the 453 phosphopeptides that showed changes with TGFβ stimulation, co-treatment with SB-525334 resulted in 203 changes in phosphopeptides, with 66 up-regulated and 137 down-regulated as compared to vehicle-treated samples (FC≥1.5, p≤0.05, Fig 3B).

**Fig 3.**
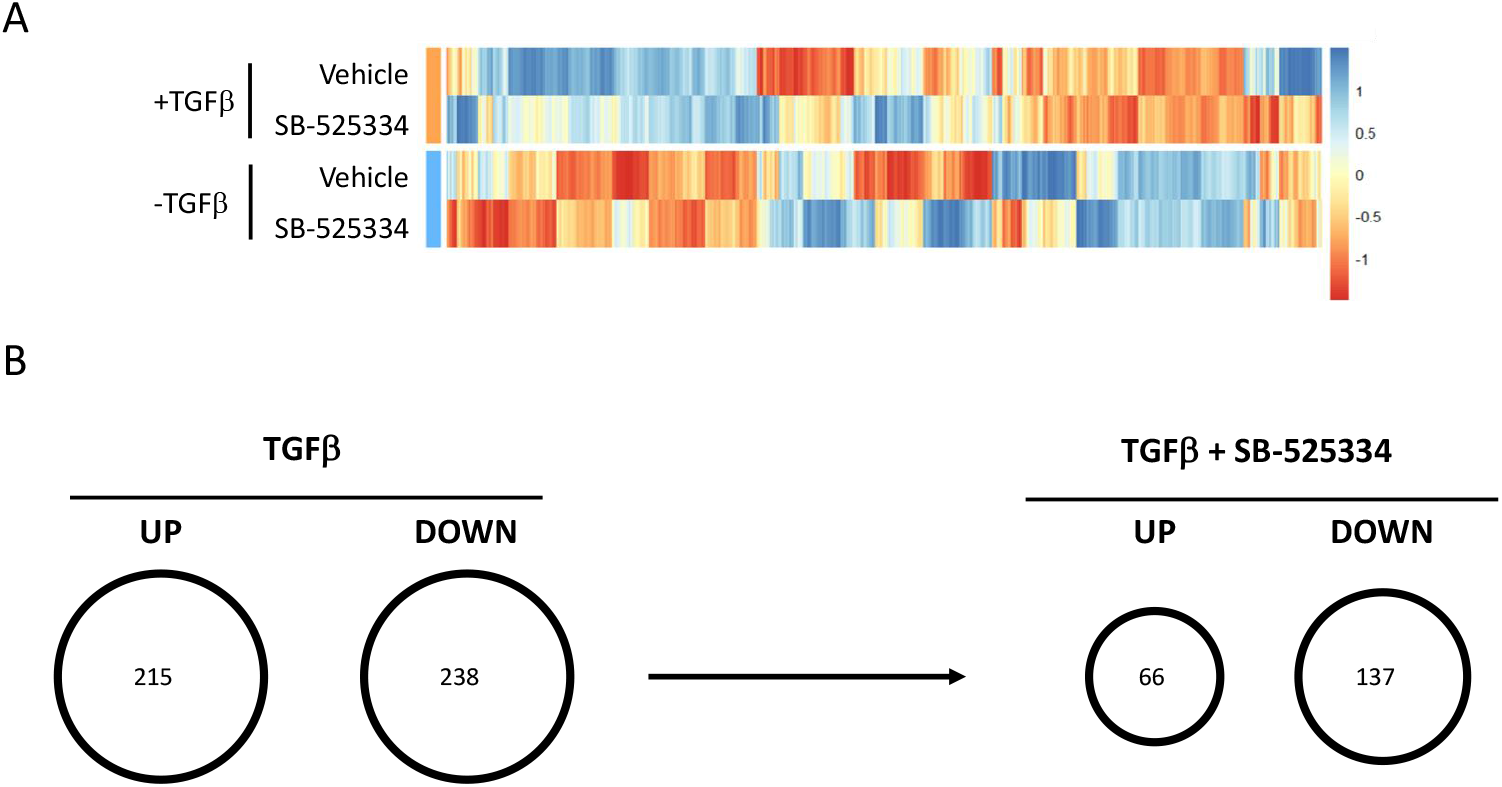
Dynamic changes of TGFβ-induced phosphorylation events by TGFβ receptor inhibitor (SB-525334). (A) Heatmap of phosphopeptides, regulated by TGFβ ± SB-525334 (color scale from blue to red indicating increased and decreased phosphorylation). (B) The Venn diagram shows phosphorylation events, significantly identified in the TGFβ ± SB-525334 treated cells (FC≥1.5 and p≤0.05 in TGFβ ± SB-525334 vs vehicle-treated groups.

Within the phosphopeptides that were changed with SB-525334 co-treatment, 37 phosphopeptides and 16 phosphopeptides showed a decrease and an increase, respectively, of at least 1.5-fold relative to TGFβ treatment alone (Tables 1 and 2).

**Table 1.** TGFβ-increased Phosphopeptides that were Antagonized by Co-treatment with SB-525334. (FC≥1.5 as compared to TGFβ treatment alone)

| Protein | Gene name | Position | Vehicle | SB-525334 | TGFβ | TGFβ<br>+SB-525334 |
| --- | --- | --- | --- | --- | --- | --- |
| Q9BWH6 | RPAP1 | Ser268 | 1.00 | 1.22 | 3.07 | 1.58 |
| Q4ADV7 | RIC1 | Ser1037 | 1.00 | 1.01 | 3.22 | 1.72 |
| P26651 | ZFP36 | Ser186 | 1.00 | 0.89 | 2.66 | 1.38 |
| P04150 | NR3C1 | Ser134; Ser45 | 1.00 | 0.94 | 2.10 | 1.09 |
| Q9NYJ8 | TAB2 | Ser450 | 1.00 | 0.98 | 2.06 | 1.06 |
| Q06190 | PPP2R3A | Ser181 | 1.00 | 0.96 | 2.44 | 1.32 |
| P25685 | DNAJB1 | Ser151 | 1.00 | 0.92 | 1.60 | 0.82 |
| Q9UPU5 | USP24 | Ser1281 | 1.00 | 1.05 | 2.29 | 1.31 |
| P98082 | DAB2 | Ser723;Thr505 | 1.00 | 0.99 | 2.08 | 1.19 |
| P04792 | HSPB1 | Ser78 | 1.00 | 0.96 | 2.01 | 1.15 |
| Q4ADV7 | RIC1 | Ser1017 | 1.00 | 0.88 | 1.83 | 1.04 |
| Q7Z7K6 | CENPV | Ser47 | 1.00 | 0.73 | 2.09 | 1.21 |
| O95816 | BAG2 | Ser20 | 1.00 | 0.84 | 1.84 | 1.08 |
| P25685 | DNAJB1 | Ser149 | 1.00 | 0.96 | 1.58 | 0.90 |
| Q9NZ32 | ACTR10 | Thr414 | 1.00 | 0.82 | 1.60 | 0.94 |
| Q9BZ29 | DOCK9 | Ser21 | 1.00 | 0.82 | 1.80 | 1.07 |
| Q15154-5 | PCM1 | Ser90 | 1.00 | 1.04 | 2.37 | 1.47 |
| Q9Y6R4-2 | MAP3K4 | Ser214 | 1.00 | 1.01 | 1.94 | 1.19 |
| Q86UU0 | BCL9L | Ser118 | 1.00 | 0.89 | 1.99 | 1.23 |
| P04792 | HSPB1 | Ser82 | 1.00 | 0.89 | 1.73 | 1.07 |
| P53667 | LIMK1 | Ser298 | 1.00 | 0.93 | 1.97 | 1.23 |
| Q4ADV7 | RIC1 | Ser1040;Ser1003 | 1.00 | 0.88 | 1.89 | 1.18 |
| A1L020 | MEX3A | Ser338 | 1.00 | 0.99 | 1.73 | 1.09 |
| P78364 | PHC1 | Ser862 | 1.00 | 0.85 | 1.77 | 1.12 |
| O15164 | TRIM24 | Ser1042 | 1.00 | 0.73 | 1.50 | 0.95 |
| Q08AD1 | CAMSAP2 | Ser464;Ser453 | 1.00 | 1.00 | 1.70 | 1.09 |
| Q13769 | THOC5 | Thr328 | 1.00 | 1.00 | 2.22 | 1.44 |
| P25054 | APC | Ser2533;Ser2432 | 1.00 | 1.03 | 2.03 | 1.32 |
| Q6KC79-2 | NIPBL | Ser1077 | 1.00 | 0.99 | 1.84 | 1.19 |
| Q9Y4G2 | PLEKHM1 | Ser482 | 1.00 | 0.82 | 1.65 | 1.07 |
| P13056 | NR2C1 | Ser64 | 1.00 | 0.92 | 1.80 | 1.17 |
| Q9UHB6-4 | LIMA1 | Ser216 | 1.00 | 0.86 | 1.76 | 1.15 |
| Q9C0A6 | SETD5 | Ser591 | 1.00 | 0.81 | 1.73 | 1.13 |
| Q15154-5 | PCM1 | Ser93 | 1.00 | 0.95 | 1.80 | 1.18 |
| O75764 | TCEA3 | Ser130 | 1.00 | 0.83 | 1.56 | 1.02 |
| O75151 | PHF2 | Ser1056 | 1.00 | 0.80 | 2.12 | 1.40 |
| P18615 | NELFE | Ser51;Thr58 | 1.00 | 0.91 | 1.64 | 1.09 |

**Table 2.** TGFβ-decreased Phosphopeptides that were Antagonized by Co-treatment with SB-525334. (FC≥1.5 as compared to TGFβ treatment alone)

| Protein | Gene name | Position | Vehicle | SB-525334 | TGFβ | TGFβ +SB-525334 |
| --- | --- | --- | --- | --- | --- | --- |
| Q14118 | DAG1 | Thr790 | 1.00 | 0.84 | 0.30 | 0.58 |
| P29317 | EPHA2 | Ser579 | 1.00 | 1.15 | 0.18 | 0.39 |
| O75781 | PALM | Ser138 | 1.00 | 1.07 | 0.46 | 0.80 |
| P60468 | SEC61B | Ser17 | 1.00 | 0.73 | 0.52 | 0.85 |
| O43194 | GPR39 | Ser396 | 1.00 | 1.03 | 0.55 | 0.89 |
| O00461 | GOLIM4 | Tyr673 | 1.00 | 0.95 | 0.47 | 0.76 |
| Q6NZI2 | PTRF | Thr279 | 1.00 | 0.75 | 0.40 | 0.64 |
| Q7Z2K8 | GPRIN1 | Ser704 | 1.00 | 1.17 | 0.50 | 0.79 |
| Q8IZD4 | DCP1B | Ser283 | 1.00 | 1.09 | 0.53 | 0.83 |
| P02786 | TFRC | Ser24 | 1.00 | 0.90 | 0.30 | 0.49 |
| P11717 | IGF2R | Ser2347 | 1.00 | 0.75 | 0.47 | 0.73 |
| Q9UN70 | PCDHGC3 | Ser733 | 1.00 | 1.16 | 0.35 | 0.55 |
| Q09666 | AHNAK | Ser3362 | 1.00 | 0.84 | 0.33 | 0.51 |
| Q9Y6M7-13 | SLC4A7 | Ser288;Ser412;Ser279;<br>Ser403 | 1.00 | 0.89 | 0.63 | 0.95 |
| Q6UVK1 | CSPG4 | 2261 | 1.00 | 0.89 | 0.36 | 0.55 |
| P43121 | MCAM | 606 | 1.00 | 0.86 | 0.27 | 0.40 |

To identify which TGFβ biological pathways were modulated under different treatment conditions, we ran IPA analyses using the enriched, selected list of significantly changed phosphopeptides (Suppl Table 1). TGFβ treatment induced phosphopeptides, associated with ERK/mitogen-activated protein kinase (MAPK), ultraviolet (UV)C-induced MAPK and integrin-linked kinase (ILK) pathways. At the same time, TGFβ treatment decreased phosphopeptides, associated with neuregulin, reelin signaling in neurons and signaling by Rho family GTPases pathways. SB-525334 significantly decreased TGFβ-induced associations in ERK/MAP, ILK, p38 MAPK and others. Some of these pathways are well-studied and -characterized as TGFβ noncanonical pathways and have optimal effects on fibrosis [24, 25]. The phosphorylation of Smad2/3 was increased in TGFβ-treated cells (Suppl Fig 2A). LC-MS was not able to quantify Smad2/3 phosphopeptides (Suppl Excel 2).

### sGC Activator Broadly Changed Phosphorylation Events in Human Lung Fibroblasts

sGCact A treatment induced changes in phosphorylation events in the human lung fibroblasts (Fig 4A). A total of 1675 phosphopeptides were either increased (n=792) or decreased (n=883) by the treatment of sGCact A alone as compared to the vehicle (FC≥1.5, p≤0.05, Fig 4B and Suppl Excel 2). Of the 1675 phosphopeptides that showed changes with sGCact A treatment alone, co-stimulation with TGFβ resulted in 1224 changes in phosphopeptides, with 602 up-regulated and 622 down-regulated as compared to the vehicle-treated samples (FC≥1.5, p≤0.05, Fig 4A). Within the 792 phosphopeptides that showed an increase with sGCact A treatment alone, TGFβ co-treatment decreased 44 phosphorylation events at least 1.5-fold (Table 3). Of the 883 phosphopeptides that showed a decrease with sGCact A treatment alone, TGFβ co-treatment increased only 3 phosphorylation events at least 1.5-fold (Table 4).

**Fig 4.**
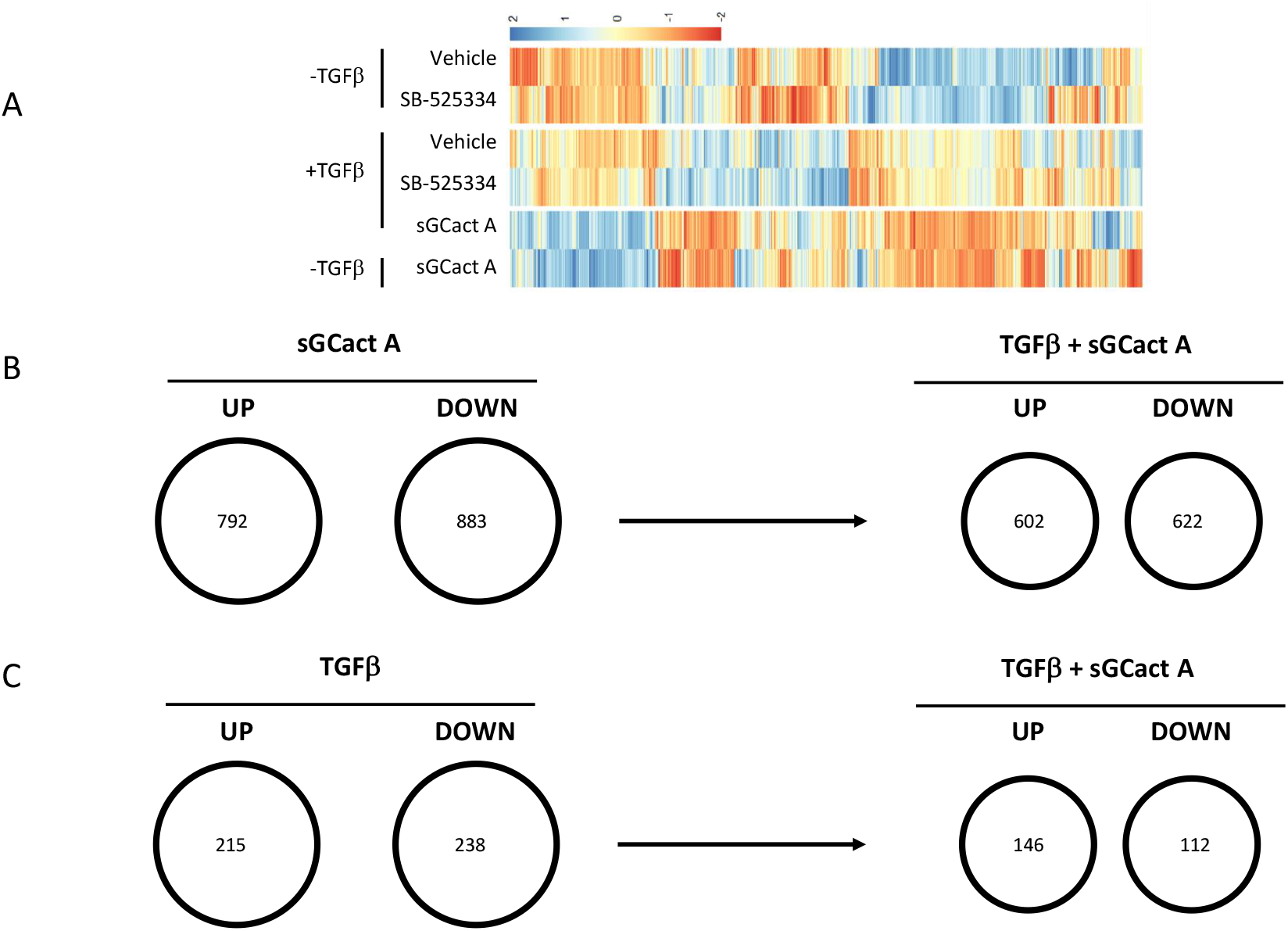
Broad changes of phosphorylation events in the sGC activator A (sGCact A)-treated human lung fibroblasts. (A) Heatmap of phosphorylation events, significantly modulated in the sGCact A ± TGFβ treated cells (color scale from blue to red indicating increased and decreased phosphorylation). (B) The Venn diagram shows phosphorylation events, identified in the sGCact A ± TGFβ treated cells (FC≥1.5 and p≤0.05 in sGCact A ± TGFβ-vs vehicle-treated groups. (C) The Ven diagram shows the effects of sGCact A on the phosphorylation events, modulated by the TGFβ treatment (FC≥1.5 and p≤0.05 in TGFβ-vs vehicle-treated groups.

**Table 3.**
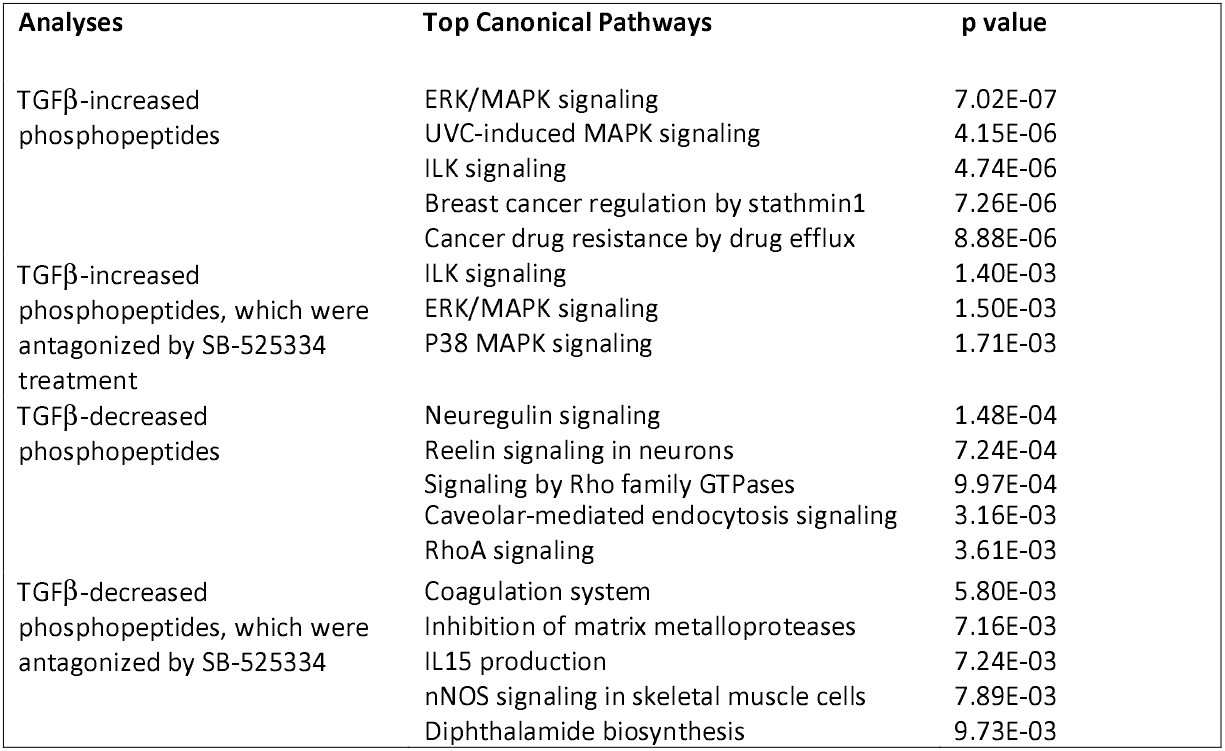
sGCact A-increased Phosphopeptides that were Antagonized by Co-treatment with TGFβ. (FC≥1.5 as compared to sGCact A treatment alone)

**Table 4.**
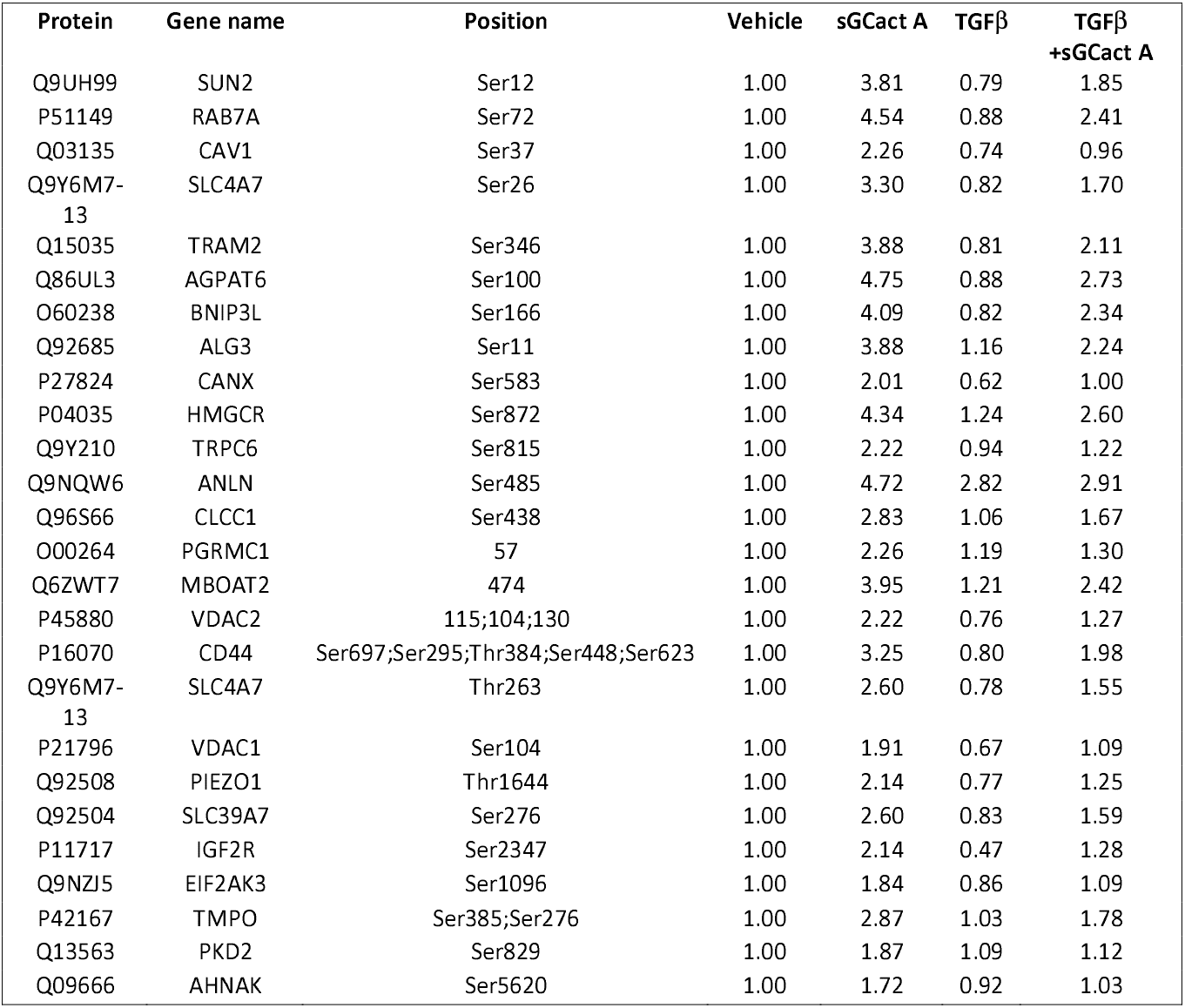
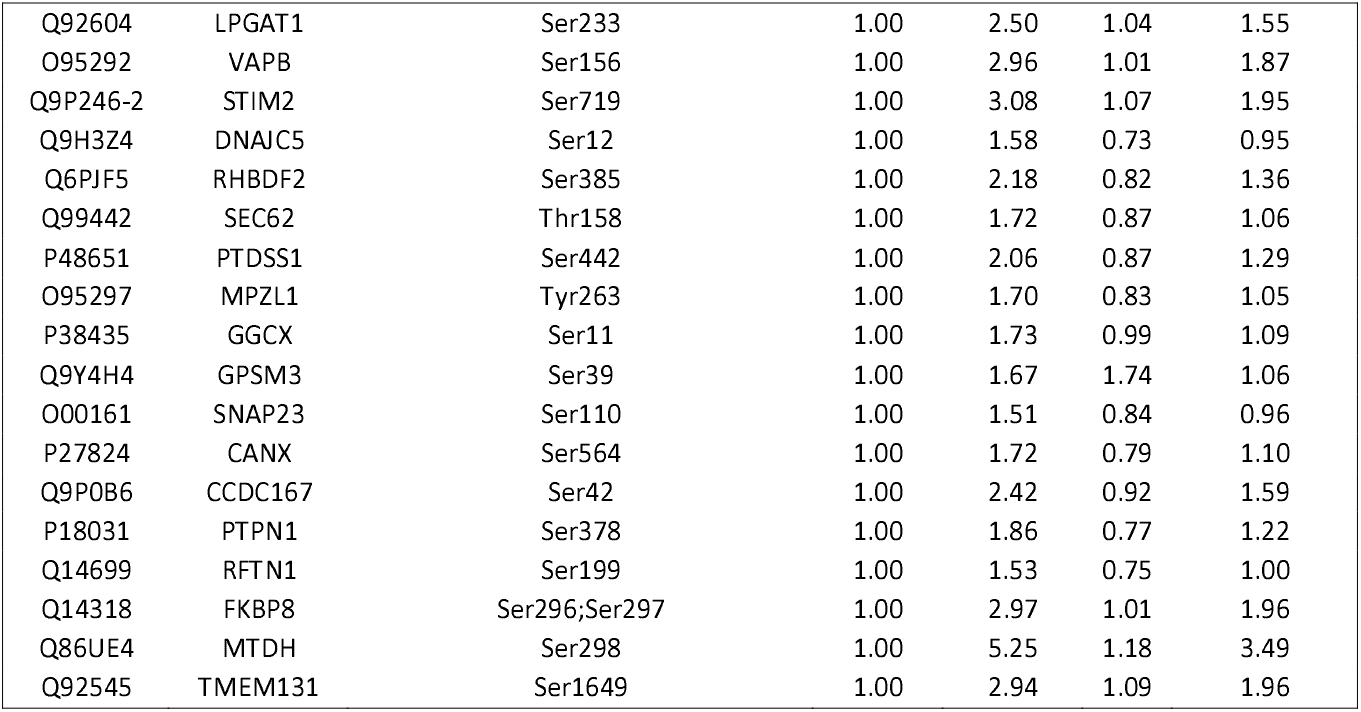
sGCact A-decreased Phosphopeptides that were Antagonized by Co-treatment with TGFβ. (FC≥1.5 as compared to sGCact A treatment alone)

To identify which biological pathways were modulated under the treatments of sGCact A, we ran IPA analyses with the list of selected phosphopeptides (Suppl Excel 2). The IPA analyses identified several affected-pathways (Table 5). sGCact A treatment induced phosphopeptides, associated with the signaling pathways of the insulin receptor, B cell receptor and gonadotropin-releasing hormone. sGCact A treatment decreased phosphopeptides associated with signaling in RhoA, GTP-binding protein Ran, protein kinase A (PKA), HIPPO, and polo-like kinase.

**Table 5.** Biological Pathways that were Associated with Phosphopeptides in sGCact A Treatment

| Protein | Gene name | Position | Vehicle | sGCact A | TGFβ | TGFβ+sGCact A |
| --- | --- | --- | --- | --- | --- | --- |
| P28290 | SSFA2 | Ser153 | 1.00 | 0.37 | 0.54 | 0.64 |
| O95391 | SLU7 | Ser515 | 1.00 | 0.34 | 0.87 | 0.60 |
| Q15424 | SAFB | Ser582 | 1.00 | 0.63 | 0.82 | 0.98 |

Of particular note, the serine/arginine repetitive matrix protein 2 (SRRM2) was identified with the highest number of unique phosphopeptides with sGCact A treatment (Table 6). This protein plays an important role in pre-mRNA splicing. A total of 27 individual Ser/Thr phosphorylation sites were significantly decreased in cells upon sGCact A treatment and this inhibition was not altered with TGFβ co-treatment. Among the 27 phosphorylation sites, 16 of these sites were previously known and reported in the UniProt database while 11 phosphorylation sites were newly identified for SRRM2 (marked with * in Table 6).

**Table 6.**
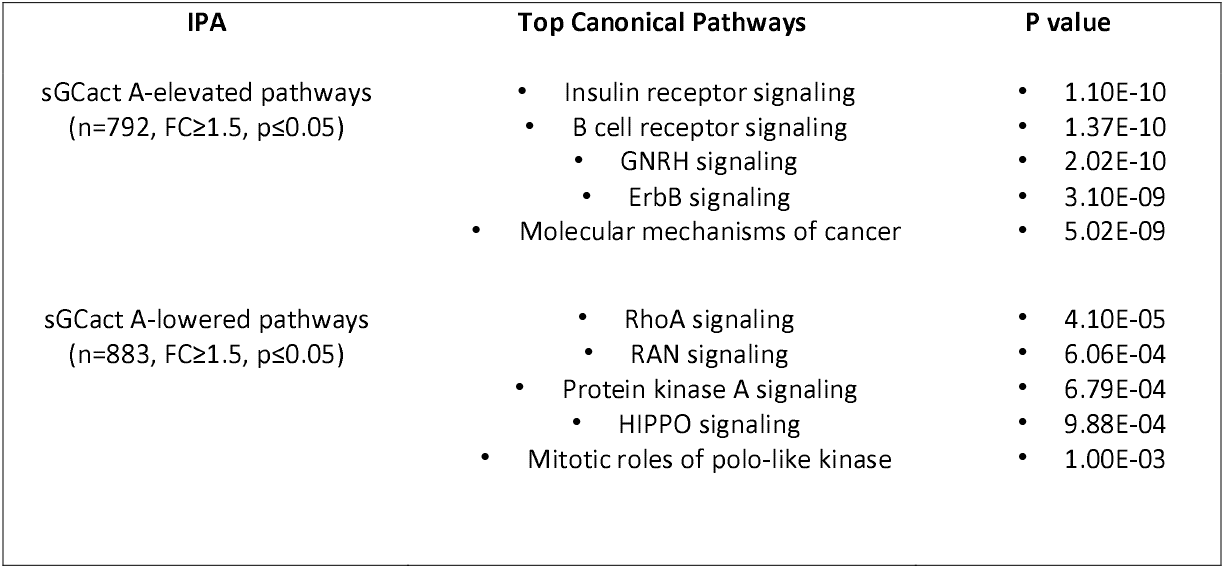
Unique Phosphorylation Changes in SRRM2

### Dynamic effect of sGC agonism on TGFβ signaling

A total of 453 phosphopeptides were either increased (n=215) or decreased (n=238) by the treatment of TGFβ alone as compared to the vehicle (FC≥1.5, p≤0.05, Fig 3B and 4C and Suppl Excel 2). Of the 453 phosphopeptides that showed changes with TGFβ stimulation, co-treatment with sGCact A resulted in 258 changes in phosphopeptides, with 146 up-regulated and 112 down-regulated as compared to the vehicle-treated samples (FC≥1.5, p≤0.05, Fig 4C).

Within the phosphopeptides that were changed with sGCact A co-treatment, 10 phosphopeptides and 52 phosphopeptides showed either a decrease or an increase, respectively, of at least 1.5-fold relative to TGFβ treatment alone (Tables 7 and 8). 11 phosphopeptides that were antagonized by sGCact A and TGFβ co-treatment compared to TGFβ treatment alone were also identified in the cells co-treated with SB-525334 and TGFβ (Table 9). The IPA analyses proposed that sGCact A modulated the association of TGFβ-induced changes in the signaling pathways of PKA, Gα12/13 and others (Table 10).

**Table 7.** TGFβ-increased Phosphopeptides that were Antagonized by Co-treatment with sGCact A. (FC≥1.5 as compared to TGFβ treatment alone)

| Protein | Gene name | Site | Vehicle | sGCactA | sGCactA+TGFβ |
| --- | --- | --- | --- | --- | --- |
| Q9UQ35 | SRRM2 | Ser1694 | 1.00 | 0.33 | 0.39 |
|  |  | Ser478* | 1.00 | 0.35 | 0.41 |
|  |  | Ser518* | 1.00 | 0.43 | 0.62 |
|  |  | Ser1693 | 1.00 | 0.44 | 0.53 |
|  |  | Ser472* | 1.00 | 0.45 | 0.50 |
|  |  | Ser783 | 1.00 | 0.46 | 0.62 |
|  |  | Ser1424 | 1.00 | 0.48 | 0.52 |
|  |  | Ser1444 | 1.00 | 0.50 | 0.59 |
|  |  | Ser1582 | 1.00 | 0.51 | 0.58 |
|  |  | Ser1436* | 1.00 | 0.52 | 0.60 |
|  |  | Ser764* | 1.00 | 0.55 | 0.69 |
|  |  | Ser456* | 1.00 | 0.56 | 0.60 |
|  |  | Ser761* | 1.00 | 0.56 | 0.70 |
|  |  | Ser231* | 1.00 | 0.57 | 0.64 |
|  |  | Ser1320 | 1.00 | 0.59 | 0.61 |
|  |  | Ser2694 | 1.00 | 0.61 | 0.62 |
|  |  | Thr1472 | 1.00 | 0.61 | 0.64 |
|  |  | Thr1492 | 1.00 | 0.61 | 0.55 |
|  |  | Ser778 | 1.00 | 0.62 | 0.66 |
|  |  | Thr1511 | 1.00 | 0.62 | 0.71 |
|  |  | Ser1857 | 1.00 | 0.63 | 0.72 |
|  |  | Ser2729* | 1.00 | 0.64 | 0.66 |
|  |  | Ser1987 | 1.00 | 0.64 | 0.75 |
|  |  | Ser820* | 1.00 | 0.64 | 0.77 |
|  |  | Ser782* | 1.00 | 0.65 | 0.66 |
|  |  | Thr2599 | 1.00 | 0.66 | 0.63 |
|  |  | Ser970 | 1.00 | 0.66 | 0.67 |

**Table 8.** TGFβ-decreased Phosphopeptides that were Antagonized by Co-treatment with sGCact A. (FC≥1.5 as compared to TGFβ treatment alone)

| Protein | Gene name | Position | Vehicle | sGCact A | TGFβ | TGFβ+sGCact A |
| --- | --- | --- | --- | --- | --- | --- |
| Q99961 | SH3GL1 | Ser288 | 1.00 | 0.74 | 1.55 | 0.61 |
| P15056 | BRAF | Ser365 | 1.00 | 1.48 | 2.11 | 1.12 |
| Q86V48 | LUZP1 | Thr958 | 1.00 | 1.00 | 1.78 | 0.94 |
| Q96EY5 | MVB12A | Ser188 | 1.00 | 0.89 | 1.58 | 0.93 |
| P50749 | RASSF2 | Thr143 | 1.00 | 0.96 | 1.72 | 1.05 |
| Q8WYP3 | RIN2 | Ser486 | 1.00 | 0.94 | 1.57 | 0.97 |
| Q96EQ0 | SGTB | Ser297 | 1.00 | 0.90 | 1.50 | 0.94 |
| Q69YQ0 | SPECC1L | Ser113 | 1.00 | 1.01 | 1.67 | 1.06 |
| Q86V48 | LUZP1 | Ser995 | 1.00 | 1.04 | 1.65 | 1.08 |
| P01106 | MYC | Ser62 | 1.00 | 1.07 | 1.59 | 1.06 |

**Table 9.** TGFβ-decreased Phosphopeptides that were Antagonized by Co-treatment with sGCact A or SB-525334. (FC≥1.5 as compared to TGFβ treatment alone)

| Protein | Gene name | Position | Vehicle | sGCact A | TGFβ | TGFβ+sGCact A |
| --- | --- | --- | --- | --- | --- | --- |
| P13639 | EEF2 | Thr57 | 1.00 | 1.99 | 0.45 | 1.75 |
| Q09666 | AHNAK | Ser5773 | 1.00 | 1.11 | 0.29 | 1.31 |
| Q5M775 | SPECC1 | Thr114 | 1.00 | 1.02 | 0.49 | 1.49 |
| P11166 | SLC2A1 | Ser226 | 1.00 | 4.44 | 0.59 | 1.59 |
| P11717 | IGF2R | Ser2347 | 1.00 | 2.14 | 0.47 | 1.28 |
| Q09666 | AHNAK | Ser5832 | 1.00 | 1.55 | 0.50 | 1.31 |
| O43194 | GPR39 | Ser396 | 1.00 | 1.32 | 0.55 | 1.30 |
| P02786 | TFRC | Ser24 | 1.00 | 2.07 | 0.30 | 0.91 |
| P05556 | ITGB1 | Thr789 | 1.00 | 2.72 | 0.64 | 1.37 |
| Q9BYG3 | NIFK | Ser145 | 1.00 | 1.19 | 0.53 | 1.19 |
| P58335-4 | ANTXR2 | Ser379;Ser276;Tyr381 | 1.00 | 1.69 | 0.46 | 1.04 |
| P43121 | MCAM | Ser606 | 1.00 | 1.87 | 0.27 | 0.72 |
| Q14118 | DAG1 | Thr790 | 1.00 | 1.39 | 0.30 | 0.74 |
| Q13443 | ADAM9 | Ser752 | 1.00 | 1.68 | 0.38 | 0.85 |
| P17302 | GJA1 | Ser306 | 1.00 | 1.92 | 0.47 | 0.97 |
| P14923 | JUP | Thr54 | 1.00 | 1.25 | 0.56 | 1.09 |
| P25116 | F2R | Ser418 | 1.00 | 1.36 | 0.48 | 0.97 |
| Q6UVK1 | CSPG4 | Thr2274 | 1.00 | 1.69 | 0.38 | 0.82 |
| P29317 | EPHA2 | Ser579 | 1.00 | 1.53 | 0.18 | 0.52 |
| Q86YV5 | SGK223 | Ser696 | 1.00 | 1.07 | 0.56 | 1.08 |
| O43493-2 | TGOLN2 | Ser70 | 1.00 | 1.56 | 0.63 | 1.15 |
| P23229-2 | ITGA6 | Tyr1059 | 1.00 | 1.86 | 0.53 | 0.98 |
| O95819-6 | MAP4K4 | Ser811 | 1.00 | 1.10 | 0.65 | 1.14 |
| P35367 | HRH1 | Ser271 | 1.00 | 1.35 | 0.39 | 0.74 |
| P51991-2 | HNRNPA3 | Ser375 | 1.00 | 0.83 | 0.40 | 0.75 |
| Q6UVK1 | CSPG4 | Thr2261 | 1.00 | 1.45 | 0.36 | 0.69 |
| O95297 | MPZL1 | Ser260 | 1.00 | 1.79 | 0.54 | 0.95 |
| Q9UBH6 | XPR1 | Ser668 | 1.00 | 1.63 | 0.50 | 0.89 |
| Q13151 | HNRNPA0 | Ser188 | 1.00 | 1.12 | 0.39 | 0.71 |
| Q6NZI2 | PTRF | Thr279 | 1.00 | 0.92 | 0.40 | 0.72 |
| Q5T036 | FAM120AOS | Ser174 | 1.00 | 1.10 | 0.41 | 0.73 |
| Q09666 | AHNAK | Ser3362 | 1.00 | 0.97 | 0.33 | 0.60 |
| Q8IUW5 | RELL1 | Ser224 | 1.00 | 1.60 | 0.48 | 0.81 |
| Q9Y250 | LZTS1 | Ser71 | 1.00 | 1.16 | 0.52 | 0.88 |
| Q14011 | CIRBP | Ser130 | 1.00 | 1.17 | 0.64 | 1.04 |
| P60468 | SEC61B | Ser17 | 1.00 | 1.16 | 0.52 | 0.85 |
| Q9UBG0 | MRC2 | Ser1453 | 1.00 | 1.15 | 0.32 | 0.56 |
| P09651 | HNRNPA1 | Ser368 | 1.00 | 0.90 | 0.41 | 0.69 |
| O15021 | MAST4 | Ser1947 | 1.00 | 0.87 | 0.52 | 0.85 |
| O43491 | EPB41L2 | Ser87 | 1.00 | 0.91 | 0.44 | 0.73 |
| Q96RR4 | CAMKK2 | Ser511 | 1.00 | 0.83 | 0.51 | 0.83 |
| Q09666 | AHNAK | Ser3360 | 1.00 | 0.98 | 0.39 | 0.64 |
| Q9C004 | SPRY4 | Ser280 | 1.00 | 1.11 | 0.44 | 0.71 |
| Q9Y5W9 | SNX11 | Ser246 | 1.00 | 1.21 | 0.60 | 0.95 |
| Q14126 | DSG2 | Ser703 | 1.00 | 1.23 | 0.48 | 0.76 |
| O15021 | MAST4 | Ser1446 | 1.00 | 0.89 | 0.66 | 1.02 |
| Q5UIP0 | RIF1 | Ser1579 | 1.00 | 0.60 | 0.46 | 0.73 |
| O00461 | GOLIM4 | Tyr673 | 1.00 | 1.16 | 0.47 | 0.74 |
| Q9BTU6 | PI4K2A | Ser468 | 1.00 | 0.98 | 0.39 | 0.59 |
| Q9H1E3 | NUCKS1 | Ser223 | 1.00 | 0.61 | 0.30 | 0.46 |
| Q9UIW2 | PLXNA1 | Ser1619 | 1.00 | 0.89 | 0.25 | 0.39 |
| Q9Y3C1 | NOP16 | Ser16 | 1.00 | 0.77 | 0.47 | 0.70 |

**Table 10.** Biological Pathways that were Associated with TGFβ-induced Changes in Phosphopeptides that were Antagonized by Co-treatment with sGCact A

**SB-525334**
| Protein | Gene name | Position | Vehicle | SB-525334 | sGCact A | TGFβ | TGFβ<br>+SB-525334 | TGFβ<br>+sGCact A |
| --- | --- | --- | --- | --- | --- | --- | --- | --- |
| Q14118 | DAG1 | Thr790 | 1.00 | 0.84 | 1.39 | 0.30 | 0.58 | 0.74 |
| P29317 | EPHA2 | Ser579 | 1.00 | 1.15 | 1.53 | 0.18 | 0.39 | 0.52 |
| P60468 | SEC61B | Ser17 | 1.00 | 0.73 | 1.16 | 0.52 | 0.85 | 0.85 |
| O43194 | GPR39 | Ser396 | 1.00 | 1.03 | 1.32 | 0.55 | 0.89 | 1.30 |
| O00461 | GOLIM4 | Tyr673 | 1.00 | 0.95 | 1.16 | 0.47 | 0.76 | 0.74 |
| Q6NZI2 | PTRF | Thr279 | 1.00 | 0.75 | 0.92 | 0.40 | 0.64 | 0.72 |
| P02786 | TFRC | Ser24 | 1.00 | 0.90 | 2.07 | 0.30 | 0.49 | 0.91 |
| P11717 | IGF2R | Ser2347 | 1.00 | 0.75 | 2.14 | 0.47 | 0.73 | 1.28 |
| Q09666 | AHNAK | Ser3362 | 1.00 | 0.84 | 0.97 | 0.33 | 0.51 | 0.60 |
| Q6UVK1 | CSPG4 | Thr2261 | 1.00 | 0.89 | 1.45 | 0.36 | 0.55 | 0.69 |
| P43121 | MCAM | Ser606 | 1.00 | 0.86 | 1.87 | 0.27 | 0.40 | 0.72 |

Our study has revealed many novel phosphorylation events, orchestrated by the sGC agonism in human lung fibroblasts as itself or through a cross-talk with TGFβ signaling. The biological implications of these novel findings have not yet been understood. Further studies with genetic and molecular approaches would be warranted.

## Discussion

The sGC enzyme exists in either a reduced or oxidized form in cells and can be pharmacologically activated by sGC stimulator or sGC activator, respectively. Under standard cell culture conditions, it can be assumed that most of the sGC pool would exist in the reduced state and thus sGC stimulator would demonstrate more pronounced effects. However, in our study using the BioMAP^®^ panels with human cells in normal culturing system, sGCact A, a selective and potent sGC activator, showed more activity relative to sGC stimulator in pulmonary fibrosis system and thus we further continued our experiments with sGCact A.

The molecular basis of sGC-cGMP agonism in vasodilation and platelet inactivation have been extensively investigated [13–15]. Several biochemical and molecular studies have revealed its downstream pathways and critical molecules and modifications in the signaling cascade to enable vascular remodeling and platelet activation. One of the most well-studied and critical molecules in this pathway is PKG, a cGMP-dependent serine/ threonine protein kinase [26, 27]. Over 1000 kinase substrates of PKG have been identified and/or proposed based on biochemical analyses, sequence motif searches and in vitro/ in vivo phosphorylation studies [26, 27]. Our study using human lung fibroblasts has also quantified many identified targets of PKG in human platelets [14, 15, 28–30], *e.g.,* ENSA-Ser^108^, protein PRRC2A-Ser^456^, ITPR3-Ser^1832^, and PDE5-Ser^102, 60^ [14, 28] (Suppl Excel 2). In addition, the totality of the phosphoproteomic data showed the most abundantly quantified phosphorylation events occurred on serine, then followed by threonine phosphorylation. These data suggest PKG activated through sGC agonism plays key roles in lung fibroblasts, analogous to what has been characterized in platelets. However, many validated phosphorylation events on the conserved PKG motif (R/K_2-3_)(X/K)(S/T)X [14] were not identified in the lung fibroblasts. For example, the known PKG-phosphorylation site of ZYX-Ser^142^ (REKV**<u>pS</u>**S) was not identified in our experiment. Instead, Ser259 of ZYX (**<u>pS</u>**P) was decreased and this site was proposed as a substrate of cyclin-dependent kinase (CDK) with a target motif of pS/T-P [31]. In support of CDK activity downstream of sGC agonism in lung fibroblasts, our data also identified increased phosphorylation events at CDK16-Thr^111^ and -Ser^184^, which correlate with its kinase activity (Suppl Excel 2). Along this notion, sGCact A-treatment antagonized TGFβ-decreased phosphorylation events on EEF2-Thr57 and SLC2A1-Ser226, which are known targets of EEF2K and protein kinase A (PKA), respectively (Suppl Excel 2). In addition to PKG and CDK, there are other serine/ threonine-specific protein kinases that play important roles in fibroblasts such as PKA, protein kinase C (PKC), MAPKs, Ca^2+^/ calmodulin-dependent protein kinases (CaMK). It is still not clear whether PKG is the dominant and major kinase protein under sGC agonism in lung fibroblasts. To better understand key molecules and essential modifications that are involved in sGC agonism-induced anti-fibrotic effects, more comprehensive molecular, biochemical and genetic studies are needed. Limited events of tyrosine phosphorylation were identified (Suppl Excel 2).

SRRM2 is a large protein with a molecular weight >300 kDa. This protein contains more than 50 serine/ threonine phosphorylation sites; however the regulation and function of these phosphorylation events are not well understood. Our study identified SRRM2 as the most broadly modified protein by the treatment of sGCact A. A total of 27 Ser/Thr phosphorylation events on SRRM2 were decreased upon sGC treatment (Table 6). Previously, two studies quantified changes in phosphorylation events of SRRM2 in the livers from simple steatosis, non-alcoholic steatohepatitis and cancer [32, 33]. In these experiments, the phosphorylation events at Thr1003, Ser1083, Thr252, Ser395 and others were increased. Among these changes, two phosphorylation events at Ser1582 and Ser1857, which were increased in liver cancer, were decreased upon sGCact A stimulation in our study. It would be interesting to address any association of these sites in cancer pathology and potential roles of sGC agonism in this process. 11 out of 27 phosphorylation events on SRRM2 were newly identified in our experiment. It would be interesting to investigate any biological implication of these newly identified modifications.

sGCact A agonism decreased TGFβ-induced phosphorylation events. However, its effect on TGFβ-increased phosphorylation events was not robust and only a total of 10 phosphorylation events showed a decrease which was more than 1.5-fold relative to TGFβ treatment alone (Table 7). The association of these proteins to fibrosis is not known. By lowering the cut-off to 1.2-fold, we identified decrease of dual specificity MAPK kinase 2 (MAP2K2)-Ser^222^ and nuclear factor kappa B p105 (NFκB1)-Ser^907^ upon sGCact A and TGFβ co-treatment relative to TGFβ treatment alone (Suppl Excel 2). ERK was previously identified as a target of sGC agonism in TGFβ-treated dermal fibroblasts [16]. MAP2K2 is a kinase protein, which phosphorylates and subsequently activates ERK [34]. Our experiment showed that sGCact A lowered TGFβ-induced phosphorylation of MAP2K2 (~20% as relative to TGFβ treatment alone) (Suppl Excel 2). This could imply that the reduction of TGFβ-induced pERK in sGC activator-treated dermal fibroblasts was due to the reduction of pMAP2K2 through sGC agonism. Also, sGCact A TGFβ cotreatment decreased the phosphorylation event at NFκB1-Ser^907^ relative to TGFβ treatment alone (~20% as relative to TGFβ treatment alone) (Suppl Excel 2). NFκB1 is an essential molecule to form the NFκB complex, which is a critical transcription factor for inflammatory responses and cell survival [35].

Interestingly but not surprisingly, LC-MS technology was not able to identify the phosphorylation events at the Smad2-Ser^465/467^ and Smad3-Ser^423/425^, which were significantly quantified by HTRF and/or by sandwich ELISA technologies (Suppl Excel 2). It is uncertain whether HTRF or sandwich ELISA has higher sensitivity than LC-MS. However, several limitations of LC-MS technology have been already discussed elsewhere [36] and raise careful caution for interpreting our data. Also in this study, we have quantified the phosphorylation events but not the expression of total proteins. Although we think there would be limited changes in each protein levels under each condition due to the short treatment time, we cannot rule out a potential impact of differentially modified protein levels on the overall phosphopeptide signals. Careful follow up studies would be warranted.

In conclusion, human lung fibroblast phosphoproteome analyses upon sGC agonism with or without TGFβ co-stimulation have provided a complex picture of sGCact A-induced changes in cellular phosphorylation events. A remarkable number of new phosphorylation sites and changes were quantified in the sGC activator-treated lung fibroblasts and described for the first time. However, the biological implication of many of these changes are still unknown. It would be important to understand how these events facilitate the anti-fibrotic efficacy of sGC agonism. Also, investigation into the biological significance of these sGCact A-induced phosphorylation events in human fibrosis would be warranted.

## Supporting information

Phosphoproteomic raw data

Selected phosphopeptides

## Abbreviations

(IPF): Idiopathic pulmonary fibrosis;
(sGC): Soluble guanylaate cyclase;
(cGMP): Cyclic guanosine monophosphate;
(sGCact): sGC activators;
(sGCstim): sGC stimulators;
(HFL1): human lung fibroblast 1;
(SMA): α-smooth muscle actin;
(SRRM2): Serine/arginine repetitive matrix protein 2

## Supporting Materials

**Suppl Fig 1. Profiling of sGC Activator and Stimulator in the BioMAP^®^ Fibrosis and Discovery Panels**.

(A) and (B) BioMAP^®^ Fibrosis and Discovery profiles for sGCact A. (C) BioMAP^®^ Fibrosis and Discovery profiles for sGC stimulator. X-axes list the quantitative protein marker readouts. Y-axes show log-transformed ratios of readouts. The grey region around the Y-axis represents the 95% significance envelope generated from historical vehicle controls. Biomarker activities are anointed when 2 or more consecutive concentrations change in the same direction relative to vehicle controls, are outside of the significance envelope, and have at least one concentration with an effect size > 20%. Antiproliferative effect is indicated by a thick grey arrow.

**Suppl Fig 2. Activation of TGFβ and sGC-cGMP Pathways.** (A) The increased phosphorylation of Smad3 upon TGFβ treatment alone was inhibited in the cells, co-treated with TGFβ and SB-525334. (B) Little to no effect of sGCact A on pSmad2/3 phosphorylation. (C) The sGCact A treatment increased VASP phosphorylation.

**Suppl Table 1. Biological Pathways that are Associated with TGFβ ± SB-525334-induced Changes**

**Suppl Excel 1. List of Protein and Peptide IDs and the Phosphopeptide Quantification Intensities of the Triplicate Samples.**

**Suppl Excel 2. Fold Changes and Significant Levels of Peptide Phosphorylation under Different Treatment Conditions.**

## Notes

### Competing Interest Statement

Sunhwa Kim, Ashmita Saigal, Weilong Zhao, Peyvand Amini, Alex M. Tamburino, Subharekha Raghavan, Saswata Talukdar are employee of Merck Sharp & Dohme Corp., a subsidiary of Merck & Co., Inc., Kenilworth, NJ, USA and stockholder at Merck & Co., Inc., Kenilworth, NJ, USA.
Maarten Hoek was an employee of Merck Sharp & Dohme Corp., a subsidiary of Merck & Co., Inc., Kenilworth, NJ, USA when the study was conducted and is currently an employee of Maze Therapeutics, South San Francisco, CA, USA. Maarten Hoek is a stockholder at Merck & Co., Inc., Kenilworth, NJ, USA.

