## Supplementary material for "Identification of the Molecular Basis of Anti-fibrotic Effects of Soluble Guanylate Cyclase Activator Using the Human Lung Fibroblast Phosphoproteome": Suppl Fig 1-2

Supplementary Figure 1. Profiling of sGCact A in the BioMAP® fibrosis and diversity plus systems

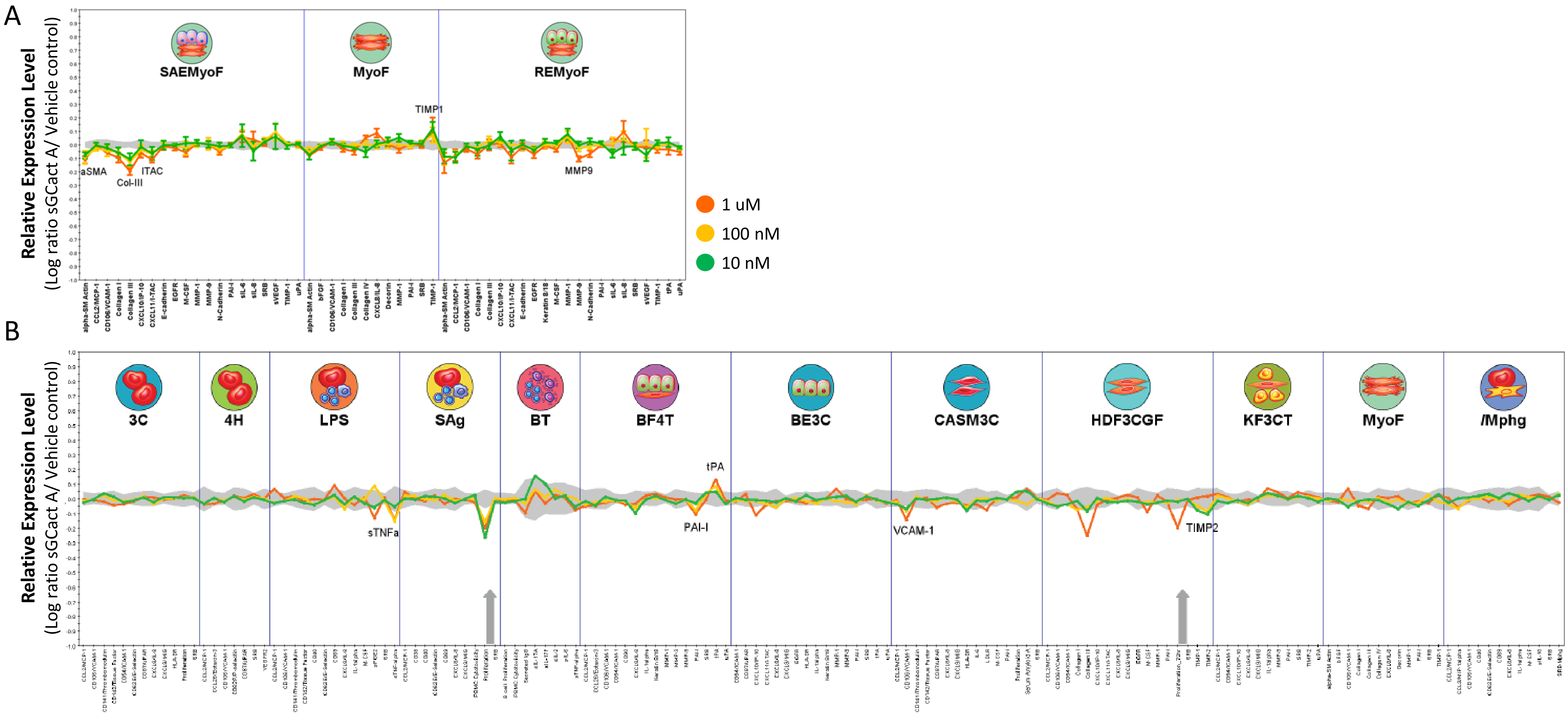

Supplementary Figure 1. continued

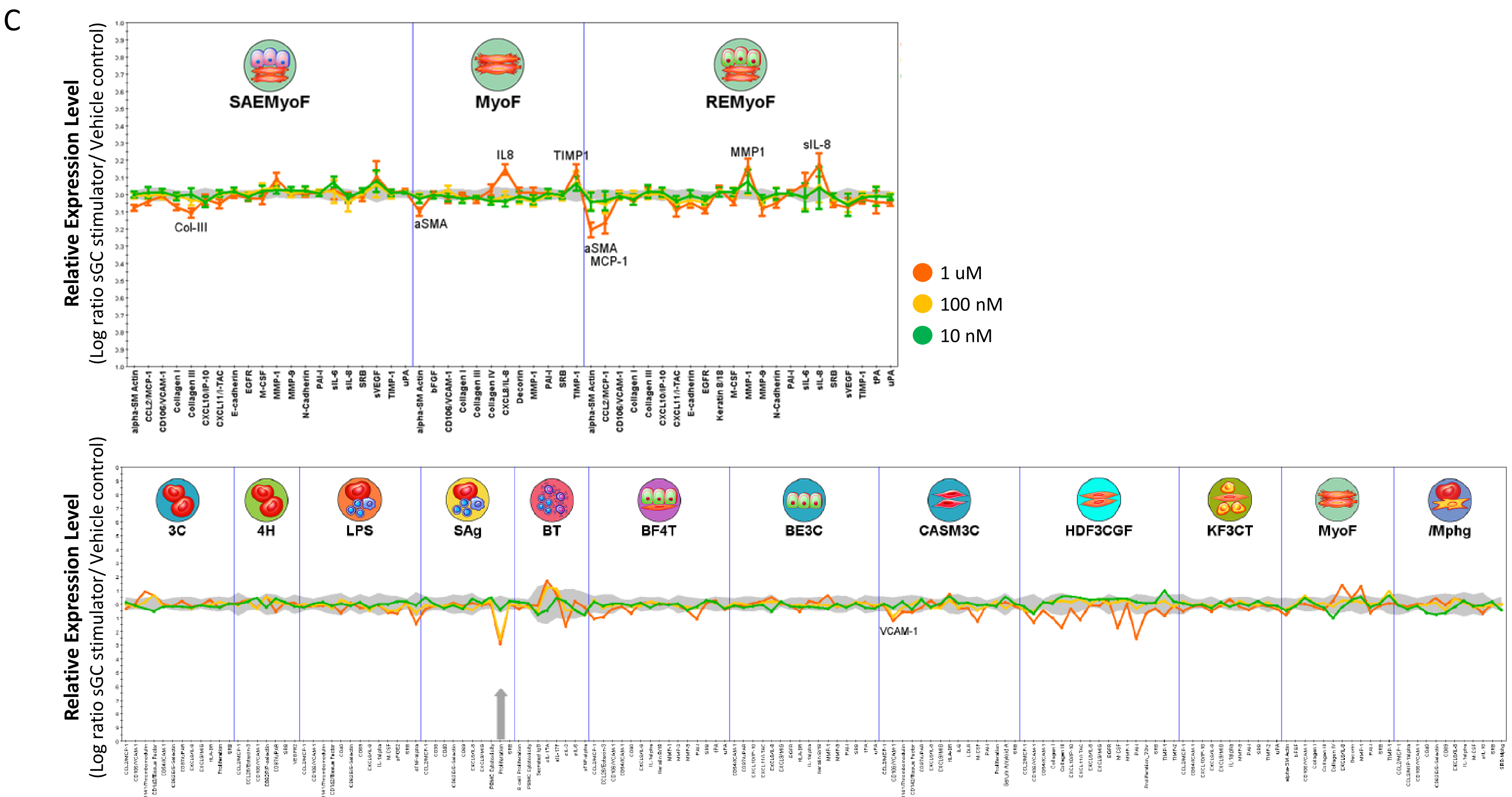

### Supplementary Figure 2. Activation of TGFβ and sGC-cGMP pathways

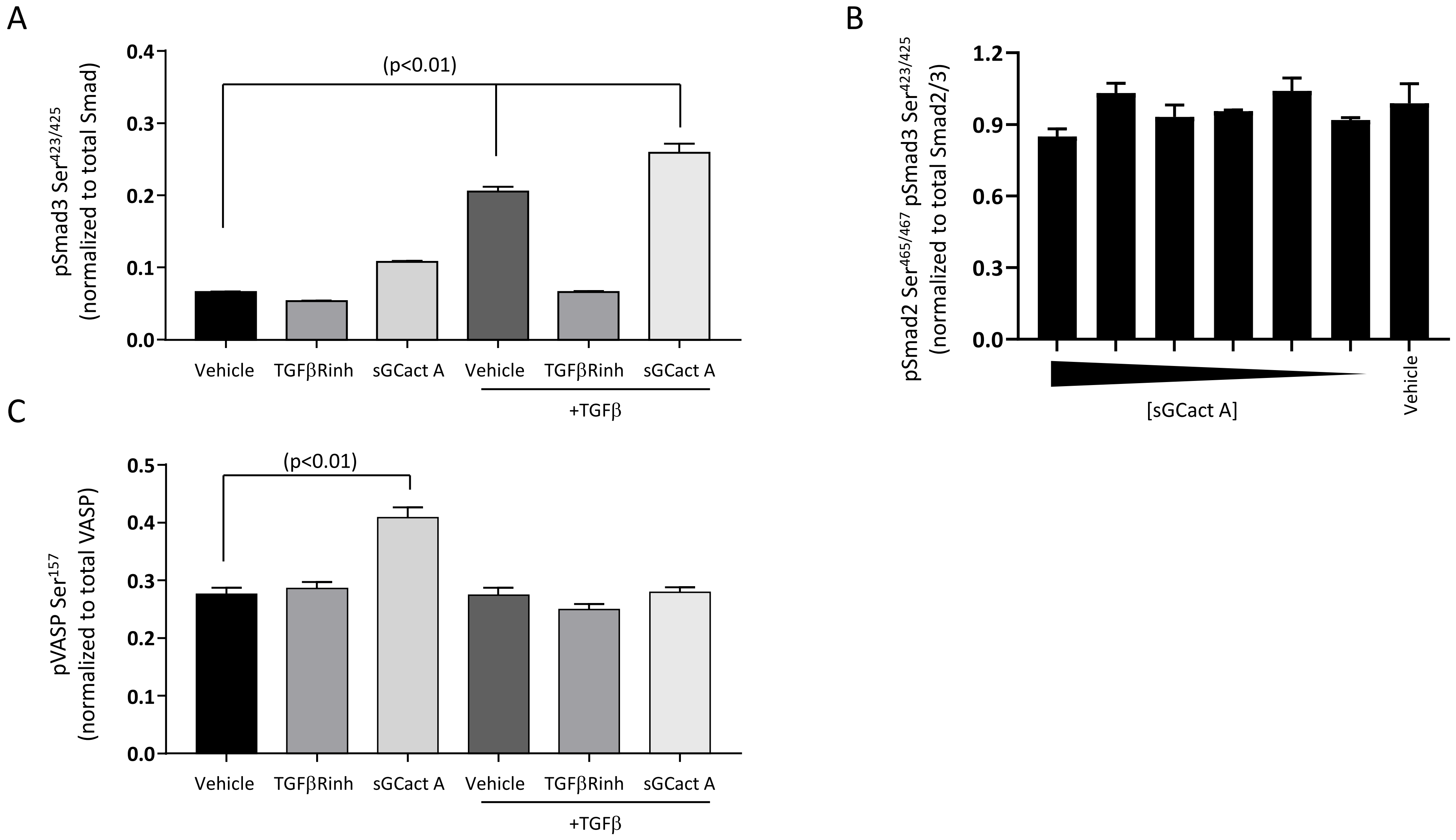
