## Supplementary material for "Identification of the Molecular Basis of Anti-fibrotic Effects of Soluble Guanylate Cyclase Activator Using the Human Lung Fibroblast Phosphoproteome": Suppl Table 1

**Supplementary Table 1. Biological Pathways that were Associated with TGFβ-induced Changes in Phosphopeptides that were Antagonized by Co-treatment with sGCact A**

| **Analyses** | **Top Canonical Pathways** | **P-Value** |
| --- | --- | --- |
| TGFβ-induced phosphopeptides that were antagonized by sGCact A treatment | Acute myeloid leukemia signaling  Protein kinase A signaling  Ga12/13 signaling  Thyroid cancer signaling  Cancer drug resistance by drug efflux | 1.83E-06  2.17E-06  8.30E-06  1.82E-05  2.69E-05 |
| TGFβ-decreased phosphopeptides that were increased by sGCact A treatment | Caveolar-mediated endocytosis signaling  Agrin interactions at neuromuscular junction  Neuregulin signaling  Granulocyte adhesion and diapedesis  Virus entry via endocytic pathways | 1.17E-04  1.58E-04  3.35E-04  3.79E-04  5.06E-04 |
